# A complete and near-perfect rhesus macaque reference genome: lessons from subtelomeric repeats and sequencing bias

**DOI:** 10.1101/2025.08.04.668424

**Authors:** Shilong Zhang, Ning Xu, Yong Lu, Yanhong Nie, Zhengtong Li, Luciana de Gennaro, Lianting Fu, Zhendong Zhang, Jieyi Chen, Kaiyue Ma, Xiangyu Yang, Juan Zhang, Matthew T. Schmitz, Francesca Antonacci, Trygve E. Bakken, Mario Ventura, Adam M. Phillippy, Qiang Sun, Yafei Mao

**Affiliations:** Bio-X Institutes, Key Laboratory for the Genetics of Developmental and Neuropsychiatric Disorders, Ministry of Education, Shanghai Jiao Tong University, Shanghai, China; Institute of Neuroscience, Center for Excellence in Brain Science and Intelligence Technology, State Key Laboratory of Neuroscience, Chinese Academy of Sciences, Shanghai, China; Shanghai Center for Brain Science and Brain-Inspired Technology, Shanghai, China; Key Laboratory of Genetic Evolution & Animal Models, Kunming Institute of Zoology, Chinese Academy of Sciences, Kunming, Yunnan, China; University of Chinese Academy of Sciences, Beijing, China; Department of Biosciences, Biotechnology and Environment, University of Bari Aldo Moro, Bari, Italy; Allen Institute for Brain Science, Seattle, WA 98109, USA; Genome Informatics Section, Center for Genomics and Data Science Research, National Human Genome Research Institute, National Institutes of Health, Bethesda, MD, USA; Center for Genomic Research, International Institutes of Medicine, Fourth Affiliated Hospital, Zhejiang University, Yiwu, Zhejiang, China; Shanghai Key Laboratory of Embryo Original Diseases, International Peace Maternity and Child Health Hospital, School of Medicine, Shanghai Jiao Tong University, Shanghai, China

## Abstract

A truly complete, telomere-to-telomere (T2T), and error-free reference genome remains a foundational resource—and long-standing goal—for unbiased comparative and functional genomics. While recent T2T assemblies of humans and other primates have made substantial progress, most still contain thousands of base-level errors, particularly within highly repetitive regions. Here, we present T2T-MMU8v2.0, a near-perfect T2T assembly of the rhesus macaque (*Macaca mulatta*), representing the highest base-level accuracy reported in a primate genome to date. By employing an optimized ONT-only assembly strategy, we identify subtelomeric satellite-rich regions as the principal bottleneck to improving assembly quality, owing to technological biases in long-read platforms and limitations in current hybrid assembly frameworks. We discover 268 previously unannotated repeat families and resolve ∼8 Mbp of SATR satellite arrays, with over 99-fold enrichment in historically misassembled subtelomeric regions. These satellites form four distinct genomic architectures, each with unique SATR satellite composition, segmental duplication organization, and epigenetic signatures, distinct from the subtelomeric architectures observed in hominid genomes. Notably, in contrast to the largely gene-poor subtelomeric regions in African hominids, the SATR architectures in macaques harbor 58 actively transcribed genes, supported by open chromatin and expression data, suggesting gene innovation within these repetitive regions. Functionally, T2T-MMU8v2.0 improves read mappability and accuracy across sequencing platforms, and results in a 19% improvement of transcription start site enrichment scores and 5,821 additional chromatin accessibility peaks on average, thereby enhancing variant detection, regulatory annotation, and transcriptomic resolution in population genetics or single-nucleus studies. Together, this work establishes a new benchmark for genomics, offers a roadmap for resolving complex repetitive regions, and reveals previously unrecognized features of subtelomeric genome structure and evolution.

## INTRODUCTION

An error-free, complete genome assembly represents the ultimate reference for unbiased comparative genomics and functional genetic analysis^1–4^. The advent of long-read sequencing technologies, combined with continual improvements in genome assembly algorithms, has enabled the generation of complete, telomere-to-telomere (T2T) assemblies^3,5–13^. Landmark accomplishments, such as the human T2T genome (T2T-CHM13) published in 2022, achieved exceptional accuracy with an estimated Phred quality value (QV) > 70, corresponding to fewer than one base error per million base pairs^6^. Similarly, the T2T macaque genome (T2T-MFA8) reached a comparable level of completeness and quality^8^. Despite these advances, current assemblies remain haploid representations and thus fail to capture the full diploid complexity of natural genomes.

The field is now advancing toward the more ambitious goal of fully phased, diploid, T2T assemblies in humans and other primates—a frontier actively being explored between 2023 and 2025^1,7,9,13^. Yet, current diploid T2T assemblies remain imperfect, often containing unplaced contigs and exhibiting lower consensus accuracy (mean QV ≈ 65) than the previous homozygous genomes (e.g., T2T-CHM13)^7,9^. At the heart of this challenge lies the difficulty of resolving extensive, highly identical repetitive regions—an issue that persists even in homozygous genomes^6,8,10^. The introduction of haplotype phasing further compounds this complexity, stretching the limits of current sequencing technologies and computational approaches^3^. Thus, recent diploid reference assemblies still contain numerous “error *k*-mers” (short sequences for which there is no underlying sequencing read support) or unexplained read mapping anomalies that suggest structural problems with the assembly.

We propose that reconstructing a complete and near-perfect haploid genome (estimated QV ≈ 100, Genome Continuity Inspector (GCI) = 100, and no unassembled/collapsed regions and unplaced contigs) represents a critical and necessary step toward the routine generation of near-perfect, complete diploid genomes. Such an assembly will resolve all previously unassembled or low-quality regions with high confidence, provide a controlled framework for identifying technical and algorithmic bottlenecks, and offer a benchmark for evaluating future assembly method performance. Insights gained from a complete and near-perfect homozygous genome will directly inform the development of future assembly tools, paving the way for accurate and automated construction of diploid and even polyploid genomes at a single-laboratory scale.

Macaques, as one of the closest genetic relatives to humans, are important nonhuman primate (NHP) models in biomedical research, underpinning progress in genomics, neuroscience, and developmental biology^14–23^. However, the current macaque reference genome (Mmul_10 or T2T-MFA8) still harbors unresolved errors^8,19^. In this study, we present a complete and near-perfect T2T haploid genome assembly for a rhesus macaque with a GCI = 100 and zero error *k*-mers remaining in the consensus sequence (*k* = 21). This assembly enabled us to address four key questions: (1) Which genomic regions were previously misassembled or missing? (2) What are the remaining technological or algorithmic barriers to achieving complete and near-perfect assemblies? (3) What biological insights can be revealed through the accurate resolution of intractable regions? (4) Does a complete and near-perfect reference improve downstream applications—such as population genetics and single-nucleus omics—compared to existing the macaque reference?

## RESULTS

### Genome assembly of a rhesus macaque

We first established a pseudo-haploid rhesus macaque (*Macaca mulatta*, MMU2019108-1) cell line that is nearly entirely homozygous diploid, as previously described^8,24^ (Supplementary Figures 1 and 2). We then generated 178 Gbp of PacBio high-fidelity (HiFi) reads, 125 Gbp of Oxford Nanopore Technology (ONT) ultra-long reads (≥ 50 kbp; N50 = 109 kbp), and 359 Gbp of Illumina short reads from this cell line (Supplementary Figure 3 and Supplementary Table 1). These datasets were used to assemble the genome with Hifiasm^12^. The resulting draft assembly spans approximately 3.09 Gbp. It exhibits high contiguity and accuracy, with a contig N50 of 146.63 Mbp, a Merqury QV^25^ estimate of 54.23, a GCI^26^ of 49.19, and a total of 55 contigs (Supplementary Tables 2 and 3).

Building on our previous T2T assembly strategy^8^, we closed 19 remaining gaps in the initial draft and reconstructed a complete rhesus macaque genome (T2T-MMU8v1.0), achieving a N50 of 162.29 Mbp and a QV of 80.11 (Supplementary Tables 4–8). These improvements are predominantly concentrated in segmental duplications (SDs) and simple satellite regions, as expected. However, read-depth–based and *k*-mer–based evaluations revealed that 9 regions and 613 error *k*-mers (*k* = 21) in T2T-MMU8v1.0, respectively, still contain misassembled sequences (Supplementary Tables 6 and 8). These error *k*-mers are predominantly enriched in subtelomeric regions (empirical *P* = 0.039, permutation test), highlighting persistent challenges in assembling subtelomeric highly repetitive sequences.

### Complete and near-perfect genome assembly and sequencing platform bias

To achieve a complete and near-perfect T2T genome assembly, it is critical to understand why certain genomic regions remain refractory to current assembly strategies (Figure 1A). We re-examined the read depth, mapping quality, and sequence characteristics of these misassembled regions. Based on these characteristics, we reassembled these regions with a custom approach and identified four major sources of bias for sequencing errors.

**Figure 1.**
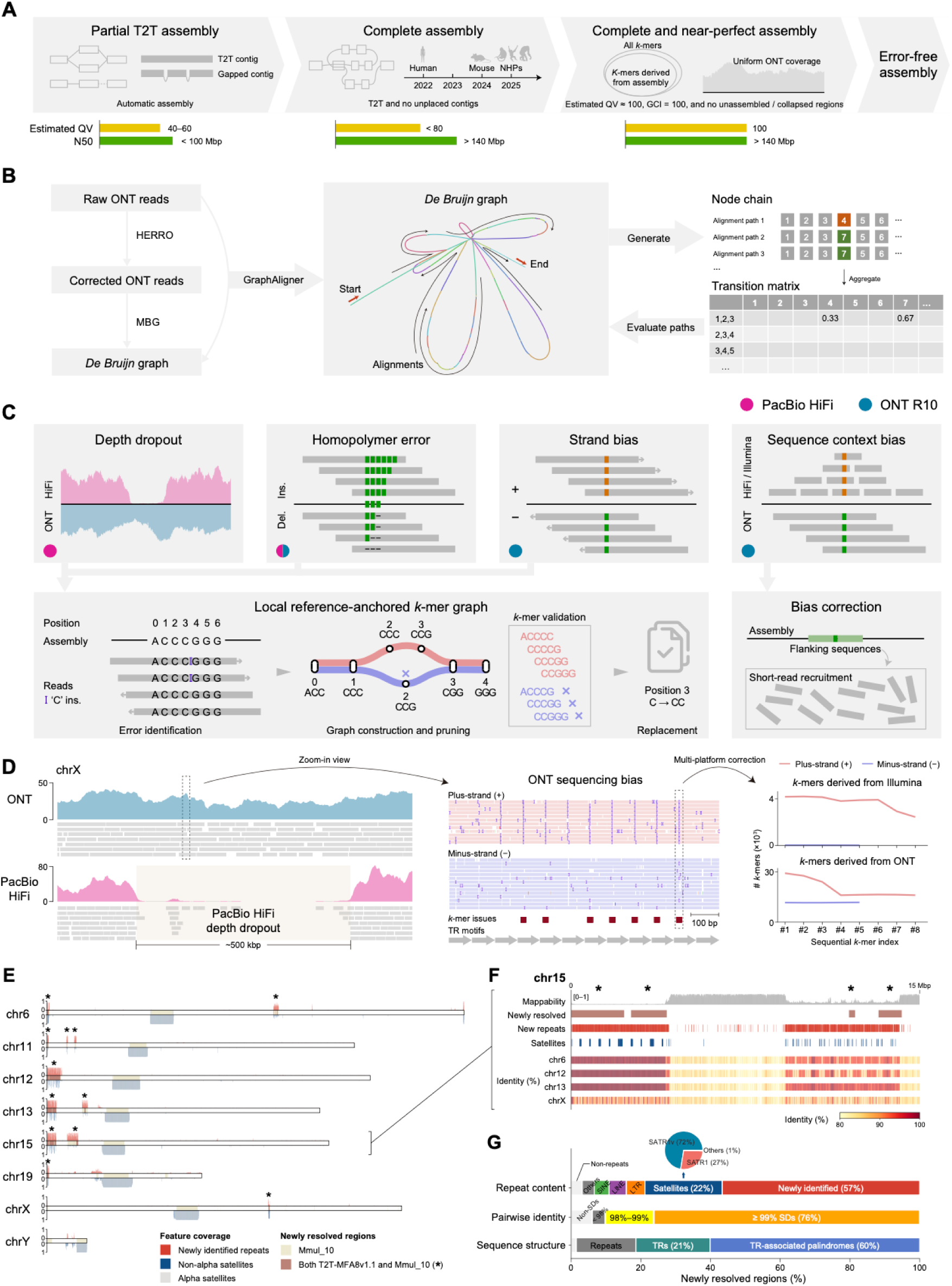
Assembly strategy and platform-specific biases in generating the T2T-MMU8v2.0 assembly. **(A)** Schematic overview of the stepwise assembly workflow, progressing from an initial draft generated by automated assemblers to the complete and near-perfect T2T-MMU8v2.0 assembly. **(B)** Custom path-finding strategy based on a corrected ONT-derived *de Bruijn* graph. The initial *de Bruijn* graph was generated corrected ONT reads with MBG, and each raw ONT read was aligned to the pruned graph. Three sequential nodes covered by a single ONT read were defined as the current state, and the fourth node as the next step. A transition matrix incorporating read-path information was then used to score enumerated candidate paths. **(C)** Representative illustrations of four types of base-pair level errors, highlighting the importance of addressing technology-driven artifacts during QV100 polishing. The schematic illustration below depicts a local reference-anchored *k*-mer graph used to resolve sequencing bias. Systematic context bias was corrected using a short-read recruitment approach. Ins, insertion; del, deletion. **(D)** Sequencing bias and polishing strategy on chrX. Left: A ∼500 kbp PacBio HiFi depth dropout region. Middle: Strand-specific ONT alignments within the dropout region reveal sequencing bias and *k*-mer distortion, with annotated TR motifs and affected *k*-mers. Right: Strand-specific *k*-mer support from Illumina and ONT reads confirms accurate homopolymer lengths in the final consensus. **(E)** The ideogram of newly resolved sequences in T2T-MFA8v2.0. Colored blocks show newly resolved sequences to Mmul_10 (tan) and those to both T2T-MFA8v1.1 and Mmul_10 (brown, indicated with asterisks). The coverage of newly identified repeats (red) and satellites (grey for alpha satellites and blue for non-alpha satellites) in 50-kbp windows are shown along the ideogram. **(F)** Repetitive landscape of newly resolved regions on chr15. Shown from top to bottom: *k*-mer mappability (*k* = 31), regions newly assembled compared to both T2T-MFA8v1.1 and Mmul_10, novel repeat families, non-alpha satellite annotations, and regions with ≥80% sequence identity to other representative chromosomes. **(G)** Composition and structure of newly resolved sequences relative to both T2T-MFA8v1.1 and Mmul_10. Bar plots show the proportions of repeat content, pairwise identity, and structural categories. The accompanying pie chart summarizes the satellite repeat family composition of these regions.

We first identified nine regions with abnormal read depth, indicative of collapses in the T2T-MMU8v1.0 assembly (Supplementary Table 8). Based on sequence content and sequence similarity relationship among the unresolved regions, we classified these into two major types: (1) four regions harboring locally high-identity tandem repeat (TR) sequences (e.g., SATR satellites on chromosome (chr) 2 and chr6), and (2) five regions with high-identity TRs across different chromosomes (e.g., SATR satellites on chr12, chr13, and chr15). These findings suggest that all unresolved regions prior to manual finishing corresponded to complex satellite TR loci.

To resolve the first class of collapses (local TRs), we constructed a minimizer-based sparse *de Bruijn* graph from corrected ONT reads^27^ using MBG^28^ as implemented in Verkko^13^, and mapped ONT reads to the local graph to trace full-length paths traversed by individual reads (Figure 1B). We then constructed a transition matrix for ONT read paths and applied a scoring function based on the Markov chain to infer the most likely ONT read path through the graph, yielding an optimized contig (Figure 1B). Using this ONT-only custom strategy, we successfully resolved four collapsed regions on chr2 and chr6 (total ∼3.5 Mbp). For the second class of collapsed regions (inter-chromosomal high-identity TRs), we first partitioned ONT reads by chromosome of origin and then combined our custom ONT-only graph-based method with Hifiasm^29^ (ONT), enabling us to resolve the collapsed subtelomeres (total ∼15 Mbp) on chromosomes 12, 13, and 15 (Methods). Following the resolution of large-scale collapses, as confirmed by uniform read depth, we focused on refining base-level errors. During polishing, we identified four distinct types of sequencing bias that underlie the remaining regions with error *k*-mers (Figure 1C).

Depth dropout is a pronounced issue in PacBio HiFi^6,13,30^, with 268 regions (∼16.68 Mbp) detected in total. By analyzing the current assembly without large-scale collapses, we found that dropout is not primarily attributable to read misalignment or misassignment, but rather reflects a true absence of HiFi reads generated by the PacBio platform (Supplementary Figure 4). This limitation likely stems from the platform’s consensus-based read generation. The affected regions are typically tandemly arranged GC-rich repeats (116 regions spanning ∼8.5 Mbp) suggesting that current chemistry and consensus algorithms struggle to meet quality criteria in such sequence contexts in the PacBio platform.

Within these dropout regions, we observe additional sequencing biases that hinder base-level accuracy. The most prominent of these are well-characterized homopolymer errors— inaccurate base calling in regions of consecutive identical nucleotides^5,30–33^ (Supplementary Figure 5). We also find strand bias, where base calling differs between plus- and minus-strand^29^, and sequence context bias, involving systematic miscalls associated with specific sequence motifs^31^ (Supplementary Figures 5 and 6). These biases may act independently or synergistically, compounding the challenge of polishing assemblies to high base-pair accuracy.

The near-homozygous nature of the cell line enables more confident resolution of these problematic regions. To address them, we employed a local reference-anchored *k*-mer graph and short-read recruitment strategy (Figure 1C; Methods). Notably, this approach revealed 190 TR loci where ONT reads consistently exhibit a GGG deletion, while Illumina-derived *k*-mers align without discrepancy (Figure 1D and Supplementary Figure 6), indicating a likely systematic ONT base-calling error in these sequence contexts. These platform-specific biases (especially within complex repetitive regions) have been underappreciated in prior studies and remain a major obstacle to achieving complete and near-perfect genome assemblies.

Following the above efforts, we generated a final genome assembly (T2T-MMU8v2.0) characterized by uniform read depth, a GCI of 100 (totally continuous alignment even with strict filtering), and a Merqury *k*-mer–based QV estimate of 100 (QV100, *k* = 21; QV77, *k* = 31) (Table 1, Supplementary Figures 7 and 8, and Supplementary Tables 9–17; Methods).

**Table 1.**
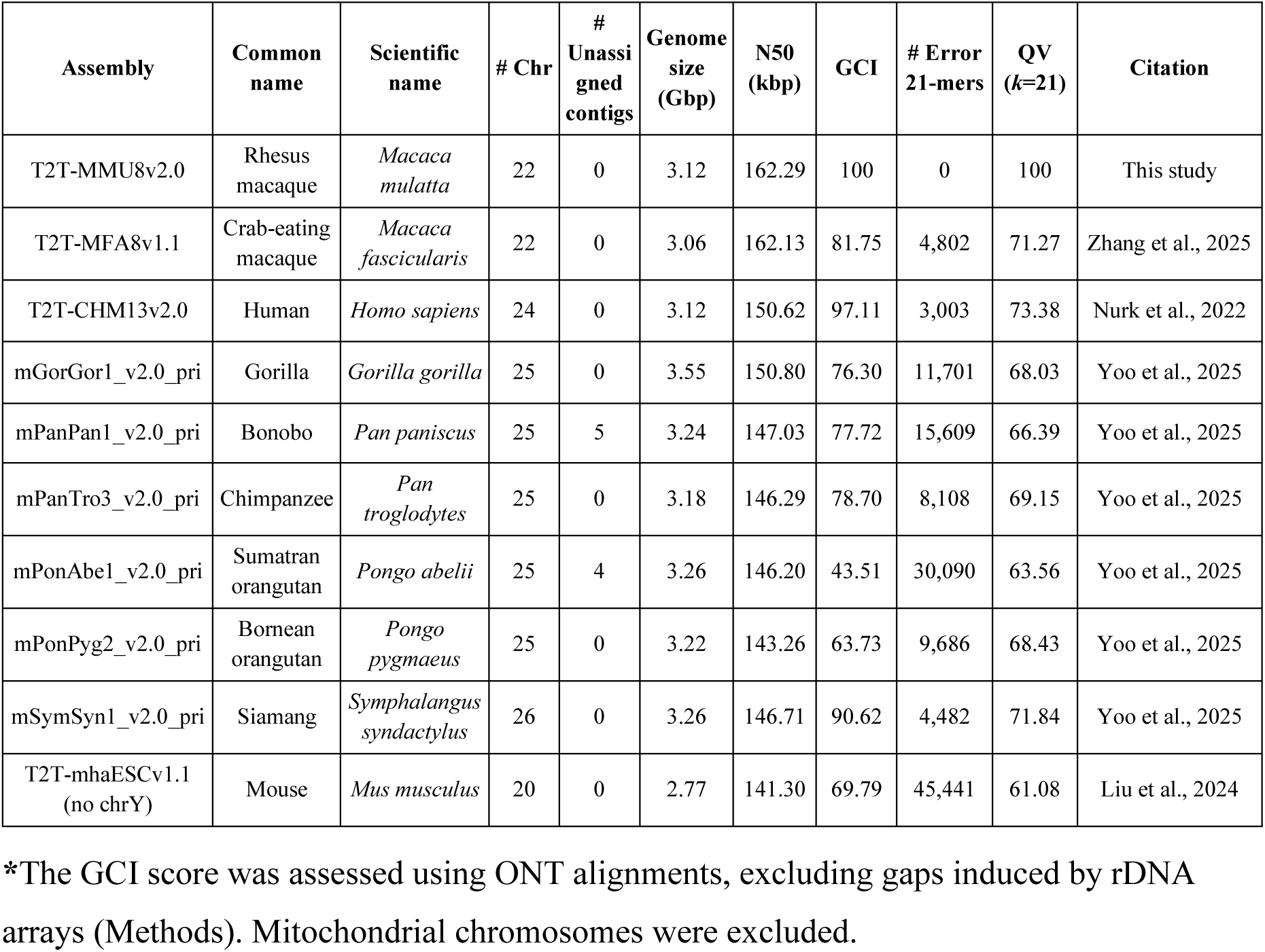
The quality evaluation of representative T2T assemblies.

| Assembly | Common name | Scientific name | # Chr | # Unassigned contigs | Genome size (Gbp) | N50 (kbp) | GCI | # Error 21-mers | QV ($k=21$) | Citation |
| --- | --- | --- | --- | --- | --- | --- | --- | --- | --- | --- |
| T2T-MMU8v2.0 | Rhesus macaque | *Macaca mulatta* | 22 | 0 | 3.12 | 162.29 | 100 | 0 | 100 | This study |
| T2T-MFA8v1.1 | Crab-eating macaque | *Macaca fascicularis* | 22 | 0 | 3.06 | 162.13 | 81.75 | 4,802 | 71.27 | Zhang et al., 2025 |
| T2T-CHM13v2.0 | Human | *Homo sapiens* | 24 | 0 | 3.12 | 150.62 | 97.11 | 3,003 | 73.38 | Nurk et al., 2022 |
| mGorGor1_v2.0_pri | Gorilla | *Gorilla gorilla* | 25 | 0 | 3.55 | 150.80 | 76.30 | 11,701 | 68.03 | Yoo et al., 2025 |
| mPanPan1_v2.0_pri | Bonobo | *Pan paniscus* | 25 | 5 | 3.24 | 147.03 | 77.72 | 15,609 | 66.39 | Yoo et al., 2025 |
| mPanTro3_v2.0_pri | Chimpanzee | *Pan troglodytes* | 25 | 0 | 3.18 | 146.29 | 78.70 | 8,108 | 69.15 | Yoo et al., 2025 |
| mPonAbe1_v2.0_pri | Sumatran orangutan | *Pongo abelii* | 25 | 4 | 3.26 | 146.20 | 43.51 | 30,090 | 63.56 | Yoo et al., 2025 |
| mPonPyg2_v2.0_pri | Bornean orangutan | *Pongo pygmaeus* | 25 | 0 | 3.22 | 143.26 | 63.73 | 9,686 | 68.43 | Yoo et al., 2025 |
| mSymSyn1_v2.0_pri | Siamang | *Symphalangus syndactylus* | 26 | 0 | 3.26 | 146.71 | 90.62 | 4,482 | 71.84 | Yoo et al., 2025 |
| T2T-mhaESCv1.1 (no chrY) | Mouse | *Mus musculus* | 20 | 0 | 2.77 | 141.30 | 69.79 | 45,441 | 61.08 | Liu et al., 2024 |

\*The GCI score was assessed using ONT alignments, excluding gaps induced by rDNA arrays (Methods). Mitochondrial chromosomes were excluded.

We resolved ∼302 Mbp and ∼25 Mbp of novel sequence absent from the Mmul_10 reference and the previously published T2T-MFA8v1.1 assembly, respectively. These regions, largely localized to subtelomeric domains (empirical *P =* 0, permutation test), are highly repetitive— comprising ∼94% SDs and ∼81% TRs with associated palindromic structures (Figures 1E–1G). Strikingly, 57% of the newly assembled sequences represent previously uncharacterized repeats^6,9,34–37^, underscoring that large portions of the subtelomeric regions remained systematically inaccessible in prior references, including complete T2T assemblies (Figure 1G). However, as expected, a genomic region composed of near-identical sequences, most notably ribosomal DNA (rDNA) arrays, remains refractory to current assembly strategies (Supplementary Figure 9). As a result, the rDNA region remains unresolved in T2T-MMU8v2.0, consistent with all previously reported T2T genomes^6,8,9^.

### Structure and epigenetic pattern of subtelomeric regions in macaques

Building on the above efforts, we found that extensive satellite TRs with palindromic configurations remain a major barrier to achieving a complete and near-perfect genome assembly. A comprehensive understanding of the genomic distribution, sequence composition, and structural organization of these satellite repeats in T2T-MMU8v2.0 may help inform generalized strategies for constructing complete and near-perfect genomes across species in the future (Supplementary Figures 10–13 and Supplementary Table 18).

Focusing on subtelomeric regions at p-arms of T2T-MMU8v2.0 (previously prone to misassembly), we observed that 18% of annotated repeats are dominated by the SATR satellite family, with the remaining 62% (*n* = 93) corresponding to previously uncharacterized repeats (total 268 in the T2T-MMU8v2.0) absent from both current primate T2T references and the Dfam database^6,9,34–37^ (Supplementary Figure 12 and Supplementary Table 18).

We identified three major SATR families, including SATR1 (3.44 Mbp), SATR1v^34^ (4.48 Mbp), and SATR2 (0.22 Mbp), which are dispersed across the entire genome but exhibit ∼99-fold enrichment in subtelomeric regions at p-arms (empirical *P* = 0, permutation test) (Figures 2A–2C and Supplementary Table 19). Specifically, 63.44% of SATR satellite arrays (5.16 Mbp) are localized to 17 subtelomeric loci, while 27.2% (2.21 Mbp) are distributed across 11 interstitial loci (Figures 2B and 2C). Notably, 99.9% of SATR1v arrays (*n* = 2,066) participate in complex mixed arrays with SATR1 or SATR2, with only two existing as standalone SATR1v arrays (Figure 2B). Similarly, around 90% of SATR1 arrays (*n* = 2,178) are found within mixed arrays. With respect to available T2T ape and human genomes, the macaque genome harbors ∼3.5-fold more SATR1 and ∼36-fold more SATR1v satellites, but ∼42% fewer SATR2 satellites^8,9^ (Figure 2A).

**Figure 2.**
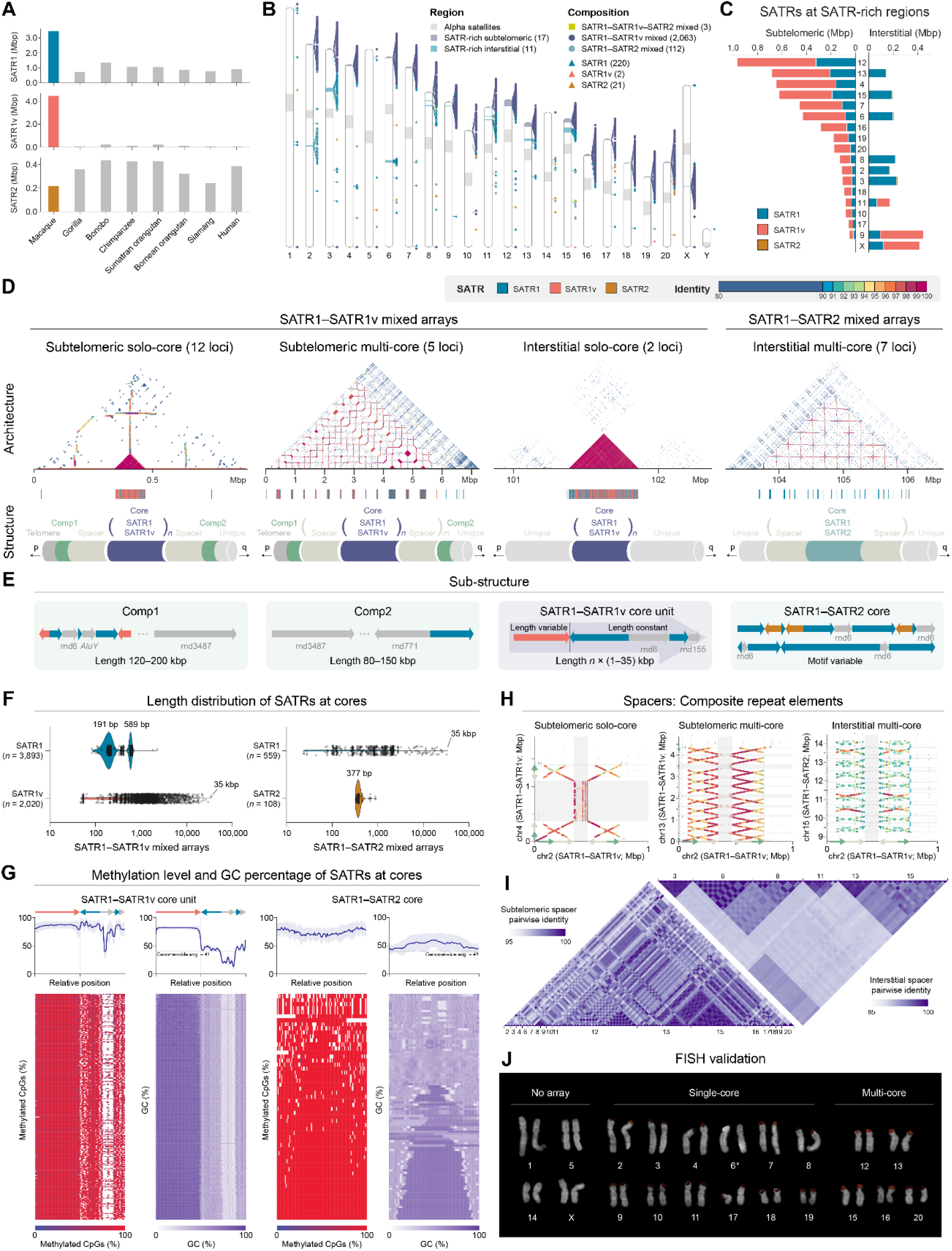
Organization, composition, and genomic distribution of SATR satellite families in T2T-MM8v2.0. **(A)** The total length of SATR1, SATR1v, and SATR2 satellite families across representative primate genomes. **(B)** Ideogram of the T2T-MMU8v2.0 with chromosomal localization of SATR arrays. Arrays composed exclusively of SATR1 (blue), SATR1v (red), or SATR2 (bronze) are indicated by triangles, respectively. Mixed arrays are shown as colored shapes: SATR1–SATR1v (dark blue circles), SATR1–SATR2 (light blue circles), and SATR1–SATR1v–SATR2 (yellow rectangles). SATR-rich genomic regions are indicated in the ideogram. **(C)** Length distribution of SATR arrays in SATR-rich regions. The left panel shows array lengths in subtelomeric regions, and the right panel shows the corresponding lengths in interstitial regions. Colors represent SATR1 (blue), SATR1v (red), and SATR2 (bronze). **(D)** Structural organization and sequence identity patterns of SATR mixed arrays. Top: heatmaps show sequence identity across four distinct array configurations — (1) subtelomeric SATR1–SATR1v solo-core mixed arrays, (2) subtelomeric SATR1– SATR1v multi-core mixed arrays, (3) interstitial SATR1–SATR1v solo-core mixed arrays, and (4) interstitial SATR1–SATR2 multi-core mixed arrays. Bottom: schematic models of the corresponding structures. Subtelomeric SATR1–SATR1v mixed arrays are flanked by conserved composite repeats and SD spacers, arranged as: telomere – left composite repeats – SD spacer – mixed arrays (core) – SD spacer – right composite repeats (from p-arm to q-arm). Interstitial SATR1–SATR1v solo-core mixed arrays lack SD spacers and flanking composite repeats. SATR1–SATR2 interstitial multiple mixed arrays are flanked by SD spacers without the aforementioned composite repeats. Telomeres (dark grey), unique sequences (light grey), composite repeats (green), SD spacers (tan), SATR1–SATR2 mixed arrays (light blue), and SATR1–SATR1v mixed arrays (dark blue) are indicated. **(E)** Repeat composition of left/right composite repeats, SATR1–SATR1v core units, and SATR1– SATR2 core units. Non-SATR repeats are shown in grey. **(F)** Length distribution of SATR1, SATR1v, and SATR2 arrays within core units. **(G)** DNA methylation and GC content profiles across core units. Top: aggregated methylation and GC percentage across individual core units. The lines represent the mean values, with shaded areas indicating mean ± s.d. Bottom: heatmaps showing methylation (blue to red) and GC content (white to purple) across individual core units. **(H)** Dot plots showing high sequence identity between SD spacers. X-axis represents composite repeats (green arrows) and SD spacers (tan arrows) and mixed arrays (grey shadow) from a representative SATR1–SATR1v mixed array; y-axis shows alignments to other mixed arrays (grey shadow) configurations with composite repeats (green arrows) and SD spacers (tan arrows) across subtelomeric and interstitial loci. Colors indicate estimated sequence identity. **(I)** Pairwise sequence identity heatmap of subtelomeric SD spacers (left; scale bar: 95%–100%) and interstitial SD spacers (right; scale bar: 85%–100%). **(J)** FISH validation of SATR arrays. Strong fluorescent signals (red) correspond to 512-bp SATR1 probes hybridizing to expanded SATR1 loci across multiple chromosomes. Chr6 (indicated by an asterisk) harbors a single SATR1 core, but barely exhibits FISH signal, suggesting potential individual polymorphism or reduced probe accessibility.

Based on the satellite composition and sequence organization of the SATR flanking regions, we identified four distinct structural architectures in T2T-MMU8v2.0 (Figure 2D, Supplementary Figures 14–22, and Supplementary Table 20). Each of these architectures contains at least one SATR core, composed of SATR arrays. Among these, three involve SATR1–SATR1v mixed arrays (*n* = 19), while the fourth consists of SATR1–SATR2 mixed arrays (*n* = 7) (Figures 2B and 2D and Supplementary Figure 16). Additionally, two loci on chr11 feature the combination of SATR1–SATR1v and SATR1–SATR2 mixed arrays. Notably, 98% and 81% of subtelomeric, and 90% and 31% of interstitial SATR architectures remained unresolved in Mmul_10 and T2T-MFA8v1.1, respectively.

Among the 19 SATR1–SATR1v loci, we further classified three structural subtypes (Figure 2D and Supplementary Table 20). At 17 subtelomeric loci, 12 exhibit a configuration with a solo core, while five display expanded multi-core structures (Figure 2D). In both cases, each locus is consistently flanked by two composite repeat elements (Figures 2D and 2E). Notably, each SATR1–SATR1v mixed array at subtelomeric loci is embedded within a stereotyped structure comprising a central SATR1–SATR1v core flanked by two spacer sequences (Figure 2D). In contrast, the other two interstitial loci harbor only the SATR1–SATR1v core and lack both spacers and composite repeats (Figure 2D and Supplementary Figure 16).

To further characterize these structures, we analyzed the sequence composition of the SATR1–SATR1v mixed arrays and their associated composite elements. Each core unit at an SATR1–SATR1v mixed array consists of one variable-length SATR1v array (GC-rich), a conserved 589-bp inverted SATR1 array, an uncharacterized repeat (rnd6; AT-rich), a second conserved 191-bp forward SATR1 array, and another uncharacterized repeat (rnd155) (Figures 2E and 2F and Supplementary Figure 23). The left-flanking composite element (comp1) at a SATR1–SATR1v mixed array spans approximately 120–200 kbp and comprises two SATR1v arrays, three SATR1 arrays, and other repeats (Figure 2F). The right-flanking composite element (comp2; 80–150 kbp) at a SATR1–SATR1v mixed array typically consists of a SATR1 array and other repeats (Figure 2F).

In contrast, the two SATR1–SATR2 mixed arrays are confined to interstitial regions and are absent from subtelomeric loci (Figure 2D). Unlike SATR1–SATR1v mixed arrays, the SATR1–SATR2 mixed arrays lack flanking composite repeats but are bordered by two spacers (Figure 2E). The motif of SATR1–SATR2 mixed cores is variable, comprising SATR1 arrays of varying lengths (ranging from 24 bp to ∼35 kbp) interspersed with uniformly sized SATR2 satellites (∼377 bp each) (Figures 2E and 2F and Supplementary Figure 23). The SATR1–SATR2 core unit is markedly more heterogeneous and less expanded than that of SATR1–SATR1v mixed arrays.

In addition to analyzing sequence content, structural organization, and array architecture, we next examined the GC content and DNA methylation profiles of the core unit in SATR mixed arrays and their spacer sequences. Core units in both SATR1–SATR1v and SATR1–SATR2 mixed arrays exhibit global hypermethylation (mean methylation level: 82% for SATR1– SATR1v core unit and 74% for SATR1–SATR2 core unit) (Figure 2G); however, they differ markedly in GC content (Figure 2G). Within the core units of the SATR1–SATR1v mixed arrays, SATR1v elements show markedly elevated GC content (mean GC = 81%), whereas SATR1 elements show a lower GC content close to the genome-wide average (mean GC = 44%) (Figure 2G). In contrast, the core units of the SATR1–SATR2 mixed arrays exhibit uniformly GC content (mean = 52%, s.d. = 0.05) across the entire array, comparable to that of SATR1–SATR1v mixed arrays (mean = 60%, s.d. = 0.23) (Figure 2G). These findings suggest that SATR1v is the primary contributor to the high GC content observed in SATR1– SATR1v mixed arrays.

Spacer sequences in subtelomeric regions (∼110–200 kbp) are highly conserved on the sequence level, with an average pairwise identity of 97.88% (ranging from 95.46% to 99.98%), whereas those in interstitial regions (∼70–230 kbp) are more divergent, averaging 91.76% identity (ranging from 88.94% to 99.73%) (Figures 2H and 2I, Supplementary Figures 24–26, and Supplementary Table 21). Notably, these spacers consistently form flanking inverted SDs at the boundaries of SATR1–SATR1v and SATR1–SATR2 mixed arrays (Figures 2H and 2I), except for interstitial spacers on chr2 (Supplementary Figures 21 and 24). This finding explains a genetic basis for the enrichment of SDs in subtelomeric regions of the macaque genome—a pattern that contrasts with humans and other great apes, in which SDs are predominantly concentrated in pericentromeric regions^8,9^ (Supplementary Figures 27 and 28). The sequence identity between subtelomeric SD spacers is ∼6% higher than that of interstitial SD spacers (*P* = 1.57×10^−17^, two-sided Mann–Whitney *U* test), which might indicate either a recent expansion or recombination between subtelomeric regions (Figure 2I). In addition, synteny analysis revealed that the SD spacers are structurally divergent, harboring large SVs within them (Supplementary Figure 29).

The structural, organizational, genetic, and epigenetic architecture of macaque subtelomeric regions is markedly distinct from the pCht arrays (satellite TRs) described in nonhuman African great apes^9,38^ (gorillas and the *Pan* lineage). The pCht arrays consist of uniform 34-bp satellite monomers and fixed-length SD spacers that display hypomethylation relative to surrounding satellite DNA^9,38^. Comparing to pCht arrays, we did not observe substantially reduced DNA methylation in macaque subtelomeric SD spacers (SATR spacer mean = 76%, pCht spacer mean = 43%) relative to the adjacent satellite arrays (SATR mean = 86%, pCht mean = 82%) (Supplementary Figure 30). These differences highlight the evolutionary divergence of subtelomeric satellite architecture and suggest lineage-specific chromatin regulation across primates.

In addition to sequence-based analyses, we performed fluorescence *in situ* hybridization (FISH) to validate the composition and chromosomal organization of SATR1 arrays. Among the 21 macaque chromosomes examined, four subtelomeric regions lacked SATR1 arrays entirely, as confirmed by FISH, while 16 exhibited clear SATR1 signals (Figure 2J). Of these, 11 harbored non-expanded arrays, whereas five showed substantial expansions. The strongest FISH signals were observed at the subtelomeres of chromosomes 12, 13, and 15, corresponding to the most highly expanded SATR1 arrays (Figure 2J). Notably, we did not observe large blocks of heterochromatin at subtelomeric regions in macaques, in contrast to the prominent subtelomeric heterochromatin observed in gorillas and the *Pan* lineage. We speculate that differences in subtelomeric structure and epigenetic regulation may underlie these cytogenetic distinctions. However, the molecular mechanisms driving the emergence, expansion, and turnover of subtelomeric regions and their biological functions in primates remain poorly understood.

Overall, the complete T2T-MMU8v2.0 assembly enables the first genome-wide fine-scale resolution of subtelomeric satellite architectures—regions that are frequently misassembled by current tools or collapsed in current macaque genomes. This high-resolution view not only offers a valuable benchmark for future algorithm development but also advances our understanding of the structural complexity in NHP genomes.

### Novel genes in the SATR satellite-associated regions

Using publicly available long-read and short-read RNA-seq data^8,39^, we annotated 21,925 protein-coding genes and 185,700 transcripts (31,423 novel) in the T2T-MMU8v2.0 assembly (Supplementary Table 22). Comparative synteny analysis with the current macaque reference genome, we identified 269 genes and 210 genes present in T2T-MMU8v2.0 but absent from Mmul_10 and T2T-MFA8v1.1, respectively (Supplementary Table 23), likely reflecting unresolved regions or incomplete annotation in the previous assemblies (Figure 3A). With respect to Mmul_10, these novel genes are ∼10-fold enriched on chrY (empirical *P* = 0, permutation test) and ∼48-fold enriched within SATR architectures (empirical *P* = 0, permutation test).

**Figure 3.**
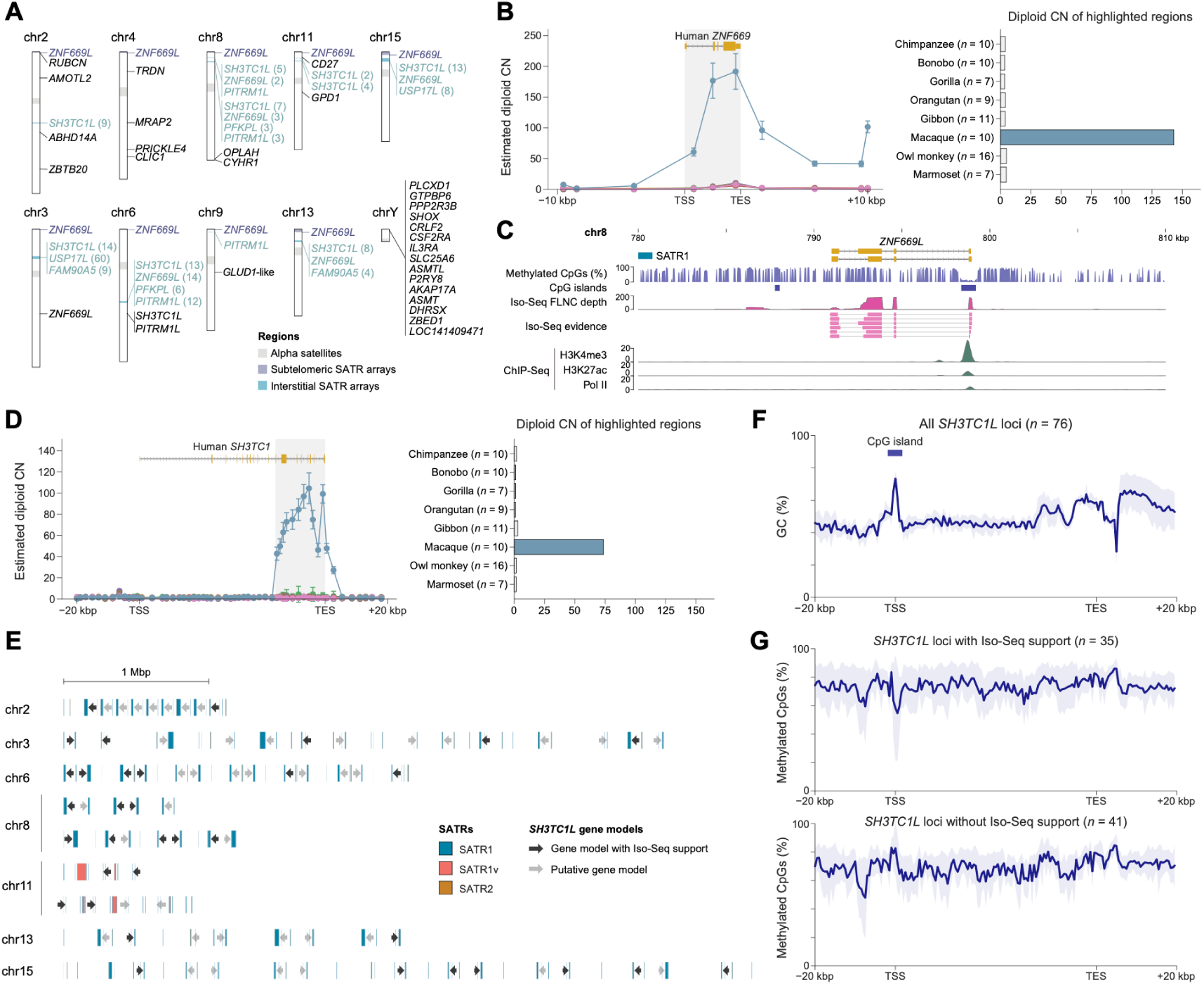
Discovery and characterization of novel genes in T2T-MMU8v2.0. **(A)** Ideograms of eight representative chromosomes from T2T-MMU8v2.0, highlighting newly annotated genes within SATR satellite-associated and previously misassembled genomic regions. **(B)** Copy number estimation at the human *ZNF669* locus shows multiple *ZNF669L* copies (left), which are absent in other primate genomes (right). **(C)** Epigenetic and transcriptomic features for a *ZNF669L* copy (∼20 kb flanking of the comp2 on chr8), including DNA methylation, Iso-Seq support, and epigenetic signals. ChIP-seq tracks are normalized by BPM (bins per million mapped reads). **(D)** Copy number estimation at the human *SH3TC1* locus reveals the expansion of truncated *SH3TC1L* copies (left), not expanded in other primates (right). **(E)** Genomic localization of *SH3TC1L* copies with (black arrows) and without (grey arrows) Iso-Seq FLNC transcript support, embedded within SATR1 (blue), SATR1v (red), and SATR2 (bronze) arrays. **(F)** GC-content profile across *SH3TC1L* loci shows well-defined CpG islands at the transcription start sites (TSSs). **(G)** Differential methylation patterns at CpG islands in *SH3TC1L* copies: transcriptionally active copies (with Iso-Seq support) show hypomethylation, while inactive copies exhibit hypermethylation, supporting the regulatory potential of active copies.

T2T-MMU8v2.0 enabled complete resolution of several previously collapsed, highly duplicated gene families corresponding to the SATR architectures. We totally identified 6 genes and 225 additional copies in these regions, including *ZNF669* (*ZNF669L*, 41 copies, 10 supported by Iso-Seq), *SH3TC1* (*SH3TC1L*, 76 copies, 35 supported), *PFKP* (*PFKPL*, 9 copies, 2 supported), *PITRM1* (*PITRM1L*, 18 copies, 11 supported), *USP17* (*USP17L*, 68 copies, none supported), as well as the primate-specific *FAM90A5*^40,41^ (13 copies, none supported) (Figure 3A and Supplementary Table 23). Among primates, *ZNF669* has undergone a lineage-specific duplication in macaques that yielded 41 truncated *ZNF669L* copies in T2T-MMU8v2.0^9,19,42^ (Figure 3B). Notably, each subtelomeric SATR architecture harbors at least one *ZNF669L* copy within the ∼10–20 kbp boundary (17 copies in total), yet only 4 exhibit both transcriptional support from full-length Iso-Seq data and epigenetic marks of active transcription (Figure 3C and Supplementary Figure 31).

In addition, we identified a previously unannotated duplicated gene, *SH3TC1*, located within SD spacers of all interstitial SATR1–SATR2 mixed arrays^43^ (Figure 3A). Similar to *ZNF669L*, the 76 newly annotated *SH3TC1L* duplicates are truncated and exhibit elevated copy number specifically in macaques relative to other primates (Figures 3D and 3E and Supplementary Table 24). All *SH3TC1L* copies (*n* = 76) harbor canonical CpG islands at their predicted transcription start sites (TSSs) (Figure 3F); however, only 46% (*n* = 35) exhibit hypomethylation at the TSSs, alongside support from full-length Iso-Seq transcripts, suggesting potential transcriptional activity and regulatory competence (Figure 3G and Supplementary Figure 32).

Together, these findings demonstrate that the T2T-MMU8v2.0 assembly not only recovers previously missing genes but also uncovers extensive, transcriptionally active duplicated copies, particularly within subtelomeric and interstitial SATR architectures. Remarkably, many of these duplicated copies (e.g., *ZNF669L*, *SH3TC1L*, and *PITRM1L*) show transcriptional activity across all 15 tissues examined, including the prefrontal cortex, spleen, and ovary (Methods), implying their function to macaque-specific traits.

### Performance improvements on read alignment and single-nucleus omics analysis

Given the enhanced resolution of previously unassembled genomic regions and gene annotation, we next evaluated the utility of the T2T-MMU8v2.0 assembly for downstream applications in population genomics and single-nucleus multi-omics.

To assess improvements in read mappability, we aligned Illumina, HiFi, ONT, and Iso-Seq full-length non-chimeric (FLNC) reads from multiple rhesus macaque individuals^8^ to both the T2T-MMU8v2.0 assembly and the current reference genome, Mmul_10. T2T-MMU8v2.0 consistently demonstrated superior mapping performance, with reduced proportions of unmapped reads, lower mismatch rates, and reduced non-primary alignments across all sequencing platforms (Figures 4A–4D). These improvements underscore the enhanced accuracy of T2T-MMU8v2.0 and its suitability as a reference for population-scale variant discovery and transcriptome analyses.

**Figure 4.**
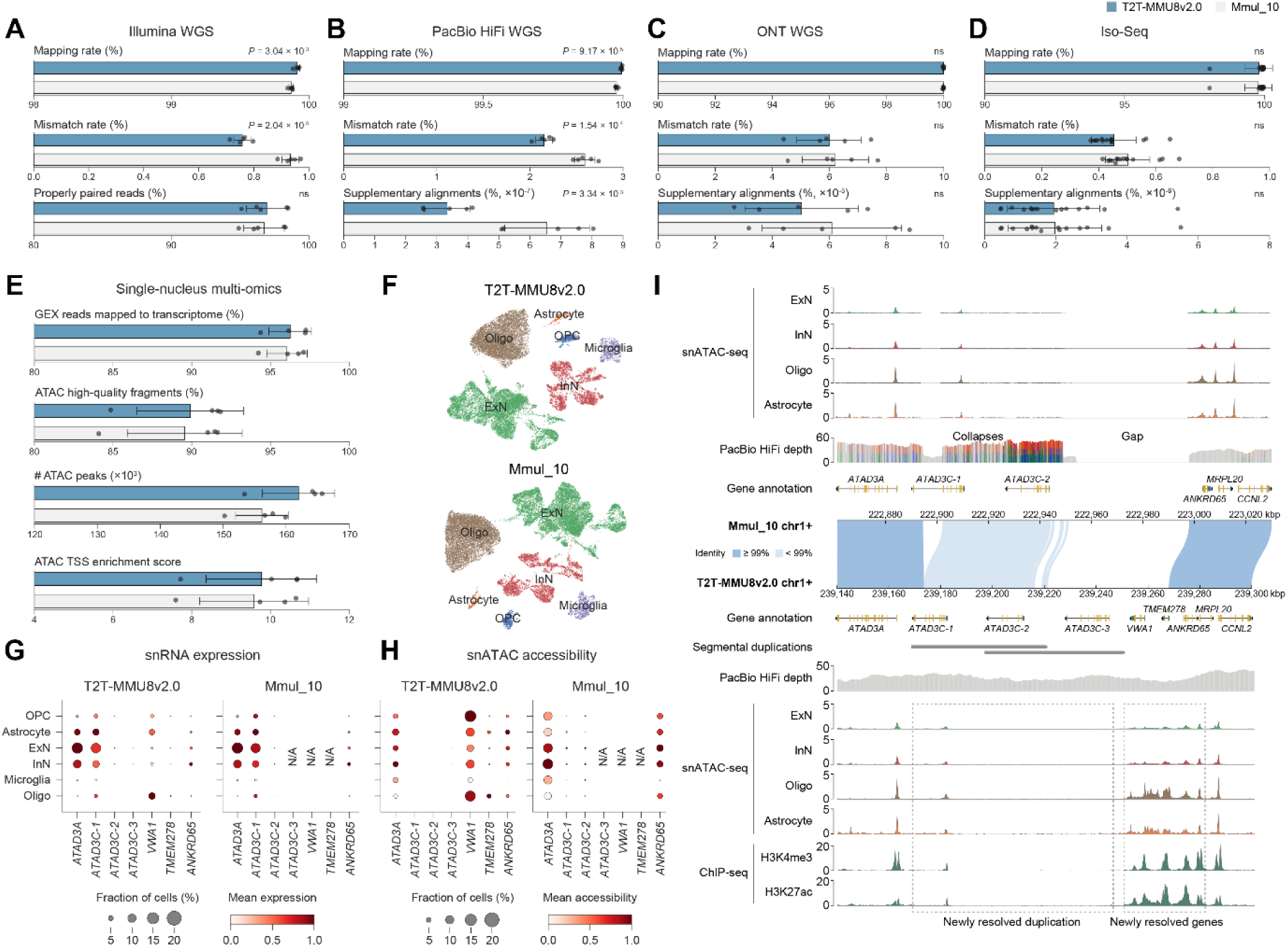
Performance comparison of T2T-MMU8v2.0 and Mmul_10 in population genetics and single-nucleus multi-omics analyses. **(A–D)** Quality control metrics for Illumina WGS (*n* = 5), PacBio HiFi WGS (*n* = 5), ONT WGS (*n* = 5), and RNA-Seq (Iso-Seq; *n* = 15) using T2T-MMU8v2.0 (blue) and Mmul_10 (grey) as references. *P* values were calculated using two-sided Welch’s *t*-tests. Error bar, mean ± s.d. ns, not significant. **(E)** Quality control metrics for single-nucleus multi-omics data (*n* = 4) using T2T-MMU8v2.0 (blue) and Mmul_10 (grey) as references. **(F)** UMAP embedding of cell types identified using T2T-MMU8v2.0 (top) and Mmul_10 (bottom) as references. **(G)** Gene expression profiles in previously misassembled regions, analyzed using snRNA-seq with T2T-MMU8v2.0 (blue) and Mmul_10 (grey) as references. **(H)** Chromatin accessibility in previously misassembled regions, analyzed using snATAC-seq with T2T-MMU8v2.0 (blue) and Mmul_10 (grey) as references. **(I)** Comparative visualization of snATAC-seq and ChIP-seq signals in previously misassembled genomic regions, using Mmul_10 (top) and T2T-MMU8v2.0 (bottom). Collapsed and gap regions in Mmul_10 are indicated by PacBio HiFi read depth. Positions with mismatch frequencies above 20% are colored, with each color representing a different substitution. Newly assembled segmental duplications are marked by grey bars, and newly resolved duplications and genes are highlighted with grey dashed boxes. snATAC-seq tracks are normalized by CPM (counts per million), while ChIP-seq tracks are normalized by BPM (bins per million mapped reads). ExN, excitatory neuron; InN, inhibitory neuron; Oligo, oligodendrocyte; OPC, oligodendrocyte progenitor cell.

To assess the impact of T2T-MMU8v2.0 on functional genomic analyses, we reanalyzed 1.04 Tbp of brain single-nucleus RNA-seq (snRNA-seq) and ATAC-seq (snATAC-seq) data^44^ (46,204 cells in total) using T2T-MMU8v2.0 and Mmul_10 as references, respectively. While snRNA-seq showed only modest gains in read mappability to the transcriptome (a 0.2% improvement), alignment to T2T-MMU8v2.0 resulted in a 0.35% increase in snATAC high-quality fragments and a 19% improvement in TSS enrichment scores, recovering an average of 5,821 additional chromatin accessibility peaks (Figure 4E). Although these differences had minimal impact on broad cell-type classification (Figure 4F), they significantly affected the detection of 20 protein-coding genes that are expressed in neurons or oligodendrocytes (Supplementary Figure 33 and Supplementary Table 25). All of these genes were misassembled in Mmul_10, resulting in their misrepresentation in expression analyses (Supplementary Figure 34). Additionally, 11,606 more chromatin accessibility peaks were identified using T2T-MMU8v2.0, highlighting the improved resolution for functional genomics analyses.

For example, the *ATAD3* locus, which is collapsed in Mmul_10, is fully resolved in T2T-MMU8v2.0, unveiling intact flanking regions where two genes—*VWA1* and *TMEM278*—are correctly assembled but entirely missing in Mmul_10 (Figures 4G–4I). Notably, *VWA1*, which encodes a von Willebrand factor A domain–containing protein with key roles in brain development^45–47^, is broadly expressed across multiple brain cell types yet was undetectable in analyses based on Mmul_10 (Figures 4G–4I). In addition, three putative regulatory elements showing chromatin accessibility in excitatory and inhibitory neurons, oligodendrocytes, and astrocytes were annotated only in T2T-MMU8v2.0 (Figures 4G and 4H). These findings highlight the critical importance of complete and accurate genome assemblies for resolving cell-type-specific gene expression and regulatory landscapes.

## DISCUSSION

An error-free, complete genome assembly represents the ultimate standard for unbiased comparative and functional genomics; yet, achieving such accuracy remains a substantial challenge^1–3^. Here, we developed a customized strategy and performed manual curation to produce a complete, near-perfect genome assembly of a rhesus macaque (T2T-MMU8v2.0). Our approach integrates QV, GCI, and read depth to generate a fully assembled genome that is devoid of error *k*-mers, unplaced contigs, and unassembled regions.

Since the release of the first complete human genome (T2T-CHM13) in 2022^6^, an increasing number of assemblies have been labeled as “complete”, “T2T”, or “gapless.”^8–10^ However, the definitions of “complete,” “error-free,” and “QV100” remain under discussion, especially given the challenges in correctly estimating assembly quality at this high level of accuracy^2,48^. Rhie et al. proposed a set of quantitative metrics to define genome quality and completeness, yet these criteria are often inconsistently applied or overlooked^25^. To our knowledge, based on five years of development and the integration of multiple T2T genome evaluation tools, the T2T-MMU8v2.0 assembly represents one of the highest standards of completeness and base-level accuracy currently achievable^6,25,26,49–53^ (Table 1).

Nevertheless, we acknowledge that these quality metrics are ultimately proxies for the true accuracy of an assembly. Despite cross-platform validation and the use of orthogonal evaluation frameworks, no current approach can fully determine absolute correctness of a genome. For example, *k*-mer–based QV estimates cannot capture all error types, such a long, homopolymer length errors or phasing errors within near-identical repeats. These limitations are exacerbated for smaller values of *k*. Therefore, we refer to this assembly as complete and near-perfect rather than error-free, emphasizing its empirical quality while acknowledging the inherent limitations of current benchmarking systems.

Based on the analysis of the T2T-MMU8v2.0 genome, our study highlights subtelomeric satellite arrays as among the most structurally complex and challenging regions to assemble. Identifying the optimal ONT read path within the initial ONT-only *de Bruijn* graph was effective for resolving these loci, though the method remains semi-automated. We propose that both this approach and the resulting genome-wide map of subtelomeric organization may inform future improvements in genome assembly algorithms. Importantly, our goal was not to develop a generalized assembler, as the scalability of this approach to non-primate or complex genomes remains unclear. Instead, our study focuses on the technological and genetic insights that can be gained from a complete, near-perfect genome, particularly in rapidly evolving, satellite-rich regions.

First, we find that the majority of error *k*-mers and unresolved sequences during the upgrade to QV100 are concentrated in subtelomeric satellite regions (Figure 1). Through targeted reassembly and polishing, we show that the ONT-only strategy offers a viable solution for resolving these regions^29^ and reveal sequence biases intrinsic to current long-read platforms (Figure 1). Second, our comparative analyses demonstrate that subtelomeric architecture differs markedly among primate species^9^, including variation in satellite array composition, structural organization, and DNA methylation patterns (Figures 2 and 3). These regions are among the most dynamic and lineage-specific in the genomes, differences that would likely remain undetected without high-quality T2T assemblies^6,8,9,34,35,38,54–56^. Our findings also expose a fundamental limitation in human-centric genomics: overreliance on the human reference genome as a benchmarking tool can obscure substantial sequence diversity and may impede the development of more broadly applicable assembly strategies. Finally, the T2T-MMU8v2.0 genome provides a robust platform for population genetics and single-cell multi-omics analysis, and serves as a high-confidence reference for comparative genomics, including cross-species structural variant discovery (Figure 4). As such, complete and near-perfect assemblies will be instrumental in advancing tool development, improving variant detection, and deepening our understanding of genome organization and evolution.

Despite these advances, several limitations remain. First, the current assembly represents a haploid genome; to our knowledge, achieving a complete and near-perfect assembly in a fully diploid genome remains an unsolved technical challenge due to the added difficulty of haplotype phasing. Second, rDNA arrays are still represented as consensus sequences in the T2T-MMU8v2.0 assembly, as current technologies are not yet able to resolve long, near-identical tandem arrays.

In summary, our study advances both sequencing methodology and biological understanding by generating a complete genome assembly with near-perfect accuracy. This work establishes a practical foundation for improved genome assembly practices and opens new avenues for investigating the structure, evolution, and function of highly repetitive genomic regions.

## Data availability

The T2T-MMU8v2.0 assembly is deposited in the National Centre for Biotechnology Information (NCBI) under accession number <u>GCA_049350105.2</u>. The raw PacBio HiFi, ONT, and Illumina WGS data of MMU2019108-1 and MMU1003063 (chrY) are deposited in the NCBI under BioProject accession number <u>PRJNA1229433</u>. T2T-MMU8 assembly, annotations, and the UCSC track hub are available at GitHub (https://github.com/zhang-shilong/T2T-MMU8) under a CC0 1.0 license. Previously published sequencing data used in this study, including <u>PRJNA269593</u>, <u>PRJNA530776</u>, <u>PRJNA602326</u>, <u>PRJNA953340</u>, <u>PRJNA976699</u>, <u>PRJNA976700</u>, <u>PRJNA976701</u>, <u>PRJNA976702</u>, <u>PRJNA986878</u>, <u>PRJNA986879</u>, <u>PRJNA1004471</u>, <u>PRJNA1037719</u>, <u>PRJNA1041301</u>, <u>PRJNA1097000</u>, and <u>PRJCA018217</u>, are available from the NCBI or the National Genomics Data Center.

## Code availability

Custom scripts used in this study are available at GitHub (https://github.com/zhang-shilong/T2T-MMU8).

## Acknowledgments

We thank Nancy F. Hansen for discussions on this manuscript. This work was supported, in part, by National Natural Science Foundation of China grants (32370658 to Y.M. and 82021001 to Q.S.); by Natural Science Foundation of Chongqing, China (CSTB2024NSCQ-JQX0004), the Computational Biology Program (24JS2840300) of Science and Technology Commission of Shanghai Municipality (STCSM), and Shanghai Jiao Tong University 2030 Initiative (WH510363003/016) to Y.M.; by National Key Research and Development Program of China (2022YFF0710901), Biological Resources Program of Chinese Academy of Sciences (KFJ-BRP-005), and National Science and Technology Innovation 2030 Major Program (2021ZD0200900) to Q.S.; and by the Intramural Research Program of the National Human Genome Research Institute, U.S. National Institutes of Health to A.M.P. The computations in this study were run on the π 2.0 cluster supported by the Center for High Performance Computing at Shanghai Jiao Tong University.

## Author contributions

Y.M. and Q.S. conceived the project. N.X., Y.L., Y.N., and Q.S. generated the cell line for sequencing and assembly. S.Z. performed all data analysis. L.d.G., F.A., and M.V. performed FISH experiments. S.Z., Z.L., L.F., Z.Z., J.C., K.M., X.Y., J.Z., M.T.S., and T.E.B. performed the genome assembly validation and application experiments. A.M.P. consulted on experimental design and validation. S.Z. and Y.M. drafted the manuscript. All authors read and approved the manuscript.

## Competing interests

The authors have declared no competing interests.

